# Sketchy understandings: Drawings reveal where students may need additional support to understand scale and abstraction in common representations of DNA

**DOI:** 10.1101/2025.02.28.640796

**Authors:** Crystal Uminski, L. Kate Wright, Dina L. Newman

## Abstract

Visual representations in molecular biology tend to follow a set of shared conventions for using certain shapes and symbols to convey information about the size and structure of nucleotides, genes, and chromosomes. Understanding how and why biologists use these conventions to represent DNA is a key part of visual literacy in molecular biology. Visual literacy, which is the ability to read and interpret visual representations, encompasses a set of skills that are necessary for biologists to effectively use models to communicate about molecular structures that cannot be directly observed. To gauge students’ visual literacy skills, we conducted semi-structured interviews with undergraduate students who had completed at least a year of biology courses. We asked students to draw and interpret figures of nucleotides, genes, and chromosomes, and we analyzed their drawings for adherence to conventions for representing scale and abstraction. We found that 77% of students made errors in representing scale and 86% of students made errors in representing abstraction. We also observed about half of the students in our sample using the conventional shapes and symbols to represent DNA in unconventional ways. These unconventional sketches may signal an incomplete understanding of the structure and function of DNA. Our findings indicate that students may need additional instructional support to interpret the conventions in common representations of DNA. We highlight opportunities for instructors to scaffold visual literacy skills into their teaching to help students better understand visual conventions for representing scale and abstraction in molecular biology.

## Introduction

Biologists rely on visual models to help them communicate about complex molecular structures that cannot be seen with the human eye. Such visual models tend to convey biology concepts through established discipline-specific visual conventions (1), and there are particular sets of shapes and symbols that biologists use to reduce complexity and to emphasize the most salient features of molecular structures for a given context (2). Consider how biologists often use a ladder shape to simply model a short DNA sequence. Conventionally, the sides of the ladder represent a sugar-phosphate backbone and the rungs of the ladder represent the nucleobase pairs. A biologist may also conventionally communicate about short DNA sequences using letters or space-filling models when the specific order of nucleotides or the chemical structure of DNA is the most relevant feature they want to emphasize. Understanding how and when to use such conventions to convey meaning is a key part of visual literacy in molecular biology (3, 4).

Like its counterpart in written communication, visual literacy in molecular biology refers to the ability to “read” and “write” using the conventions of discipline-specific symbols and notations that encode meaning in visual representations (1, 5). Experts in molecular biology visual literacy can accurately “read” or decode how conventions are used in a visual model to convey pertinent information and can “write” or draw a representation using conventions to communicate biology concepts to others (6). Developing expertise in molecular biology visual literacy is an integral part of molecular biology education. Visual literacy is so central to molecular biology education that the skills associated with “reading” and “writing” using visual models are encompassed within the *Vision and Change* core competencies for “Modeling” and “Communication and Collaboration” (7, 8).

Despite the importance of visual literacy in molecular biology, our previous work indicated that undergraduate students may need additional support to develop expert-level visual literacy for molecular biology concepts (9, 10). Developing visual literacy may be particularly challenging for students because biologists can represent molecular structures at different scales and often use different abstract symbols to emphasize varying aspects of the same structure (1, 5, 11, 12). Previous research hypothesized that challenges in understanding the conventions of scale and abstraction in visual models may be primary constraints on the development of visual literacy (6, 13).

The DNA Landscape (12) is a useful conceptual framework for delineating the conventions of how biologists represent DNA at different levels of scale and abstraction. The DNA Landscape was developed after reviewing how DNA was represented across thousands of textbook figures, and the resulting conceptual framework is presented as a three-by-three matrix. Representations of DNA can be mapped to a location on the DNA Landscape based on scale (nucleotide, gene, chromosome) and the degree of abstraction (literal shape, elements of shape and abstraction, very abstract). The DNA Landscape emerged from an analysis of textbooks that were written and edited by biology experts, so the way that scale and abstraction is portrayed in the DNA Landscape reflects the conventions accepted and used by biology experts.

We can use the DNA Landscape framework to identify erroneous or unconventional ways that biology students are “writing” with the conventions that biologists use to represent DNA. If a student’s sketch (or the verbal description of what their sketch represents) does not follow the conventions typically associated with a specific location on the DNA Landscape, this signals that the student may not correctly understand how and why biologists use certain symbols to represent DNA. We anticipate that misuse or misunderstanding of the typical visual conventions for representing DNA is an indicator that the student likely holds deeper conceptual errors about DNA structure or function.

Proficiency in molecular biology visual literacy relies on a foundational content knowledge in biology as well as an understanding of how biologists conventionally represent such content in diagrams and figures. Here, we designed an interview protocol to specifically probe the content knowledge and visual literacy skills of undergraduate students, in which students were asked to sketch representations of DNA. We analyzed student sketches generated during the interview protocol to answer the following research question: “In what ways are students misusing or misunderstanding the common conventions for representing scale and abstraction in molecular biology?”

## Methods

We conducted 45-minute semi-structured interviews with 35 students from two institutions (Table 1). All research participants had completed at least a year of undergraduate biology coursework. We recruited participants via institutional email, flyers, in-person recruitment during upper-level biology courses, and snowball sampling. During the recruitment process, we notified participants that they would be completing sketches during the interview and we asked them to arrive to the interview prepared with a pen and paper. We compensated participants with a $20 Amazon gift card. This research was approved and classified as exempt from human-subjects review by Rochester Institute of Technology (protocol 01090823).

**Table 1.** Research participant demographics.

| Table 1. Research participant demographics |  |  |  |
| --- | --- | --- | --- |
|  | **PUI** | **R2** | **%** |
| *Gender* |  |  |  |
| Man | 3 | 3 | 17 |
| Woman | 8 | 18 | 74 |
| Non-binary or gender queer | 1 | 2 | 9 |
| *Year in college* |  |  |  |
| Sophomore | 4 | 9 | 37 |
| Junior | 5 | 6 | 31 |
| Senior | 3 | 5 | 23 |
| Master's or Continuing Education | 0 | 3 | 9 |

Our interview protocol involved a series of tasks designed to elicit specific molecular biology visual literacy skills across levels of Bloom’s Taxonomy (14, 15). The protocol was composed of three parts which followed the same series of tasks that were modified in each part to focus on a different topic in molecular biology (chromosomes, nucleotides, genes). We present a generalized and a specific set of interview tasks in Table 2.

**Table 2.** Tasks completed by interview participants and alignment of interview tasks to levels of Bloom’s Taxonomy.

|  |  |  |
| --- | --- | --- |
| Generalized interview task | Specific interview prompt | Targeted Bloom's Taxonomy level(s) |
| Provide a verbal definition or description | "What is a gene?" | Remember |
| Make a sketch | "Can you sketch a gene?" | Remember/Understand |
| Explain the sketch | "Why did you draw the gene that way?" | Remember/Understand |
| Explain the figure from a concept assessment item | "Explain what you think about this representation of a gene." | Remember/Understand |
| Evaluate the strengths and limitations of the figure | "What about this image makes it an effective representation of a gene?" | Evaluate |
| Compare the sketch to the figure | "How does your sketch of a gene compare to this representation?" | Analyze |
| Explain or draw what it would look like to "zoom into" this molecular structure | "Please sketch or describe what it would look like if you "zoomed in" to a gene." | Remember/Understand |
| Explain or draw what it would look like to "zoom out" of this molecular structure | "Please sketch or describe what it would look like if you "zoomed out" of a gene." | Remember/Understand |
| Select the answer to a concept assessment item | "Please talk through how you answered this question." | Apply |
| Translate the answer from the concept assessment item into a new sketch | "Can you sketch a representation of the answer you selected?" | Apply |
| Note: Bloom's Taxonomy levels classified based on the Visualization Blooming Tool (15) |  |  |

Within the interview protocol, we asked participants to explain figures from published concept assessment instruments when the figures were presented in isolation, and then asked them to answer the entire concept assessment item in a later task during the interview. The concept assessment items were Item #2 from the Genetics Concept Assessment (16), Item #10067 from the BioMolViz assessment library (17), and Item #38 from GenBio-MAPS (18), which included an abstract model of unreplicated chromosomes, a space-filling model of DNA nucleotides, and a box-and-line representation of a gene, respectively. We used these items because they each included a conventional representation of a molecular biology structure and the items had been vetted through rigorous assessment validation procedures by the developers of each assessment.

We conducted and recorded interviews using Zoom. Of the 35 interviews, 11 participants created drawings using the Zoom whiteboard and 24 participants drew on paper then subsequently sent us photos or scans of their sketches via email. The quality of the drawings and sketches on the Zoom whiteboards may have varied depending on the familiarity and comfort with the platform.

Students generated 277 unique sketches of nucleotides, genes, and chromosomes during the interviews. We coded each sketch for alignment to the DNA Landscape (12). Using the visual conventions within the DNA Landscape as a reference, we used structural coding (19) to code each sketch for erroneous or unconventional uses of scale and abstraction (Table 3). We considered sketches in the context of how students described what they drew. In some cases, participants drew what appeared to be an unconventional representation but verbally acknowledged and clarified the ways in which their sketch differed from their mental image of what they were trying to represent. When the participant followed their sketch with a verbal description of how they would amend their sketch to better align with typical conventions, we considered these sketches to be conventional.

**Table 3.** Codes for analyzing scale and abstraction in student sketches.

| Table 3. Codes for analyzing scale and abstraction in student sketches |  |
| --- | --- |
| Code | Code description |
| Erroneous scale | The participant used or described symbols in their sketch that reflected a factual error in their understanding of the size of a particular molecular structure or the scalar relationship between multiple molecular structures. |
| Erroneous abstraction | The participant used or described symbols in their sketch that reflected a misunderstanding of the visual conventions that are typically used to represent a particular molecular structure or concept. |
| Unconventional scale | The participant used or described symbols in their sketch to convey factually correct information about the size of a particular molecular structure or the scalar relationship between multiple molecular structures, but their use or description of symbols differ from how an expert might use symbols to convey the same information. |
| Unconventional abstraction | The participant used or described symbols in their sketch to correctly convey factually correct information about a particular molecular structure or concept, but their use or description of symbols differ from how an expert might use symbols to convey the same information. |

To establish interrater reliability, we randomly selected 10% of the sketches in the sample (n = 27) and two raters (CU and DLN) independently reviewed the sketches and associated transcript for alignment to the DNA Landscape and for the use of erroneous or unconventional scale and abstraction. The two raters met to discuss any disagreements. We refined our codebook based on where disagreements occurred. The two raters repeated this process with a second set of sketches (n = 27). After establishing interrater reliability in independent coding, one rater coded the remainder of sketches in the sample. Both raters subsequently met to review the application of codes to the entire sample, and consensus values for each sketch are reflected in the final dataset.

## Results

We found that the majority of students think about the commonly-used shapes and symbols in molecular biology representations in different ways than experts might. Across the 35 students we interviewed, 97% (n = 34) drew at least one sketch that represented nucleotides, genes, or chromosomes, in ways that were misaligned to the conventional ways that experts typically represent the same structures. Here, we identify the types of molecular structures students drew and the frequency of the misuses or misunderstandings of the common conventions for representing scale and abstraction in molecular biology. We provide exemplar sketches that illustrate the ways that students were using shapes and symbols in ways that suggest an incomplete understanding of foundational molecular biology concepts.

### What did students draw?

We identified 277 student sketches that aligned to locations within the DNA Landscape (Figure 1). The majority of sketches were “highly abstract” representations consisting of letters or simple shapes. Sketches often contained multiple elements from across the DNA Landscape. These findings are not necessarily surprising, as experts often communicate with abstract representations across scales. The higher incidences of chromosome representations may reflect that chromosomes were the first subject in the sequence of the interview protocol and that many students drew chromosome-scale structures in their depictions of genes (e.g., representing a gene as a chunk of a chromosome arm).

**Figure 1.**
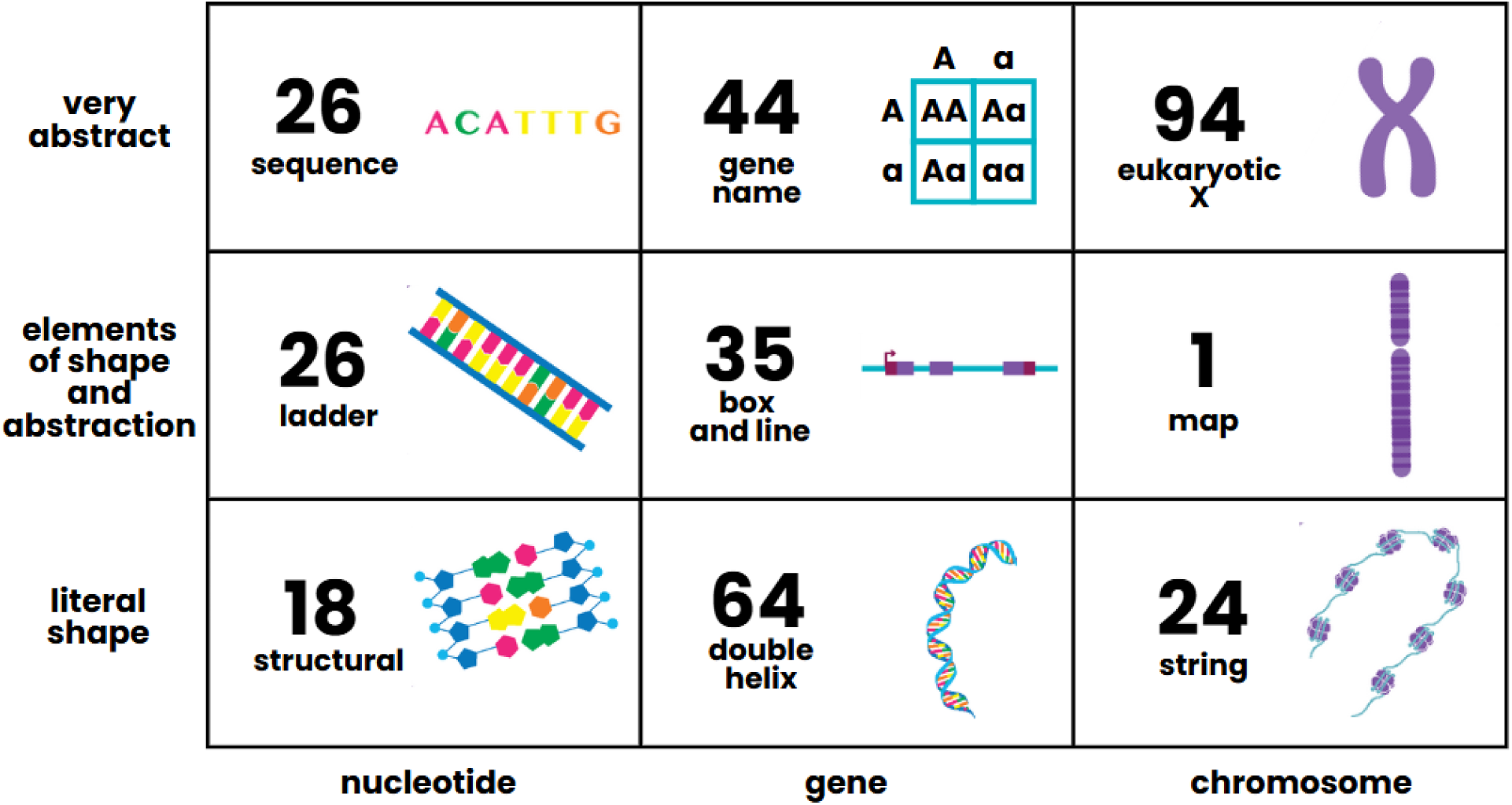
Number of student sketches aligned to each location on the DNA Landscape. Sketches that contained elements from multiple locations on the DNA Landscape were coded in each location, so the sum of sketches here exceeds the 277 total sketches generated in the interviews.

### How often did sketches contain errors or unconventional representations of scale and abstraction?

Overall, approximately half of the sketches in our sample (49%, n = 137) reflected erroneous or unconventional depictions of scale and/or abstraction to represent structures in molecular biology (Table 4). While only about a third of sketches contained errors in abstraction (29%), we found that 86% of students drew at least one sketch with an abstraction error. We similarly found that even though the total number of sketches with scale errors was relatively small (17% of sketches), over three-quarters of students made a scale error in at least one sketch. Unconventional abstraction was less common, comprising only 10% of the sketches. Notably, we did not identify any instances of unconventional scale.

**Table 4.** Errors or unconventional uses of scale and abstraction in student sketches.

|  |  |  |  |  |
| --- | --- | --- | --- | --- |
| Code | Number of sketches | Percent of sketches | Number of participants | Percent of participants |
| Scale error | 47 | 17% | 27 | 77% |
| Abstraction error | 80 | 29% | 30 | 86% |
| Unconventional scale | 0 | 0% | 0 | 0% |
| Unconventional abstraction | 27 | 10% | 17 | 49% |
| Note: These counts include a total of 14 sketches that had co-occurrences of errors in scale and errors in abstraction and 3 sketches that had co-occurrences of errors in scale and unconventional abstraction. |  |  |  |  |

We found that there were associations between the location of a sketch on the DNA Landscape and the frequency of errors in scale or abstraction. The “highly abstract” representations in the the DNA Landscape (nucleotide sequence, gene name, Chromosome X) often had higher frequencies of abstraction errors (Figure 2). While abstraction errors occurred across nearly all locations in the DNA Landscape, there was a clear pattern that conventions of “highly abstract” representations were a frequent source of misunderstanding for students. We also identified student sketches with scale errors across many of the DNA Landscape locations, with the most frequent scale errors occurring in sketches of nucleotide ladders and gene helices.

**Figure 2.**
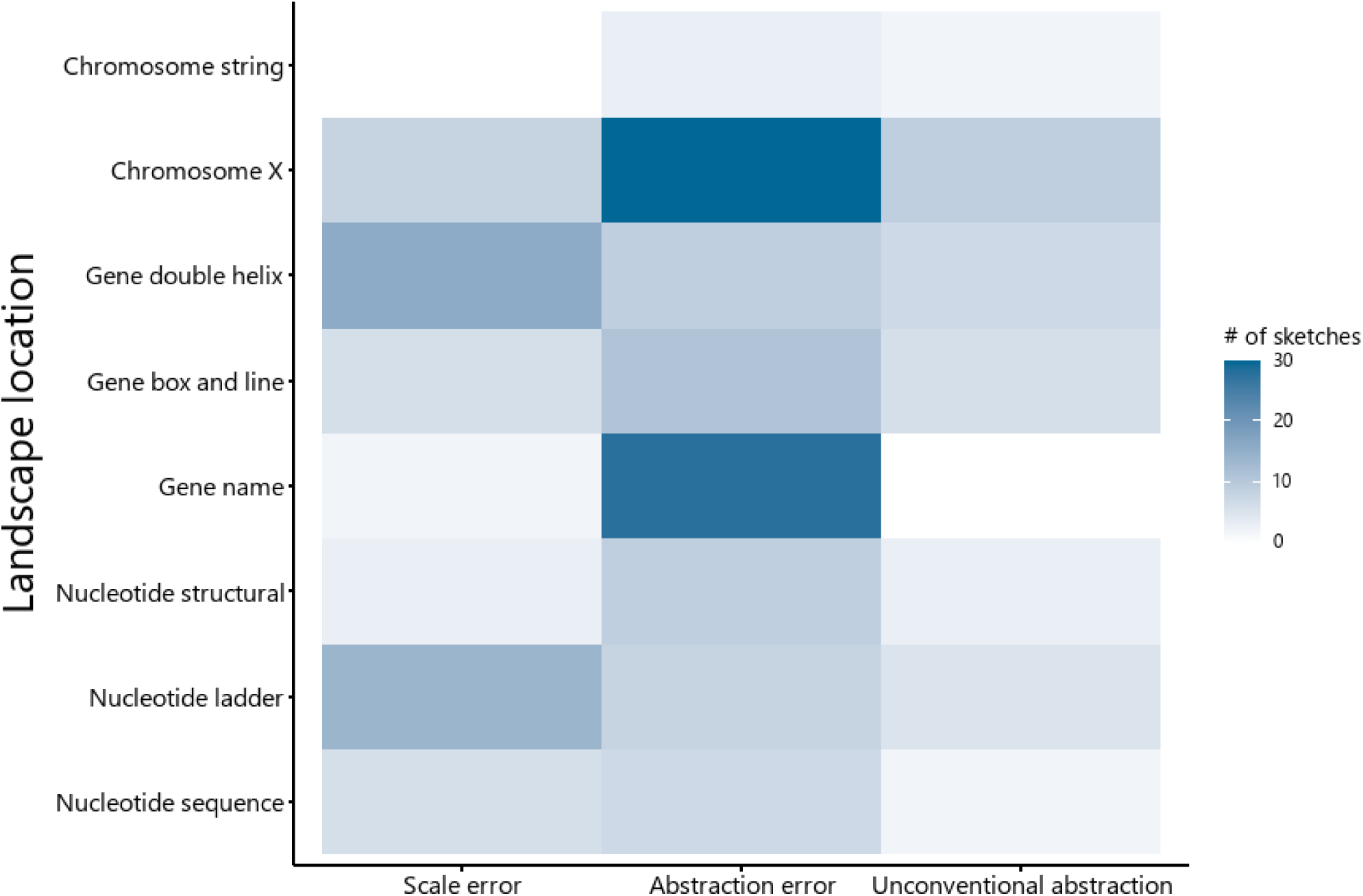
Heat map indicating the frequency of scale errors, abstraction errors, or unconventional abstraction in sketches aligned to locations on the DNA Landscape. Sketches with errors in scale reflect a factual error in their understanding of the size of a particular molecular structure or the scalar relationship between multiple molecular structures. Sketches with errors in abstraction reflect a misunderstanding of the visual conventions that are typically used to represent a particular molecular structure or concept. Sketches with unconventional abstraction convey factually correct information about the size of a particular molecular structure or the scalar relationship between multiple molecular structures, but their use or description of symbols differ from how an expert might use symbols to convey the same information. Sketches that contained elements from multiple locations on the DNA Landscape were coded in each location, so the sum of sketches here exceeds the 277 total sketches generated in the interviews. There were no errors in the single sketch of a chromosome map, so this data point is omitted here.

### What did errors in scale look like?

We provide a few illustrative examples of errors in representing scale in Figure 3. Many of the errors in scale reflected a misunderstanding the conventions for representing the size of genes. Consider how both Alex and Donna used the helical and ladder representations of DNA, respectively, to indicate that the length of a gene is approximately 3 nucleotides. We see additional misuses of conventional gene symbolism in Alex’s sketch in which the start and stop codons are boxes approximately the same size as the boxes for introns and exons in a typical box-and-line representation. We want to emphasize the mismatch between the sketches that Alex and Donna drew and the words that they used to describe their sketches. Alex demonstrates an understanding that a gene is a “section of the DNA that codes for your proteins,” and Donna similarly states that “a gene is a sequence of DNA,” yet the sketches each student drew indicate that they are substantially underestimating just how “big” the section or sequence of DNA is in a typical gene. While Alex and Donna underestimated the size of a gene, we saw Sheryl making overestimates. Sheryl drew a single gene, labeled as *SRY* in her sketch, which comprises nearly half the length of an arm of a chromosome. All the examples in Figure 3 notably contain elements from multiple locations on the DNA Landscape, which was a common, but not defining, characteristic of many of the sketches that contained errors in scale.

**Figure 3.**
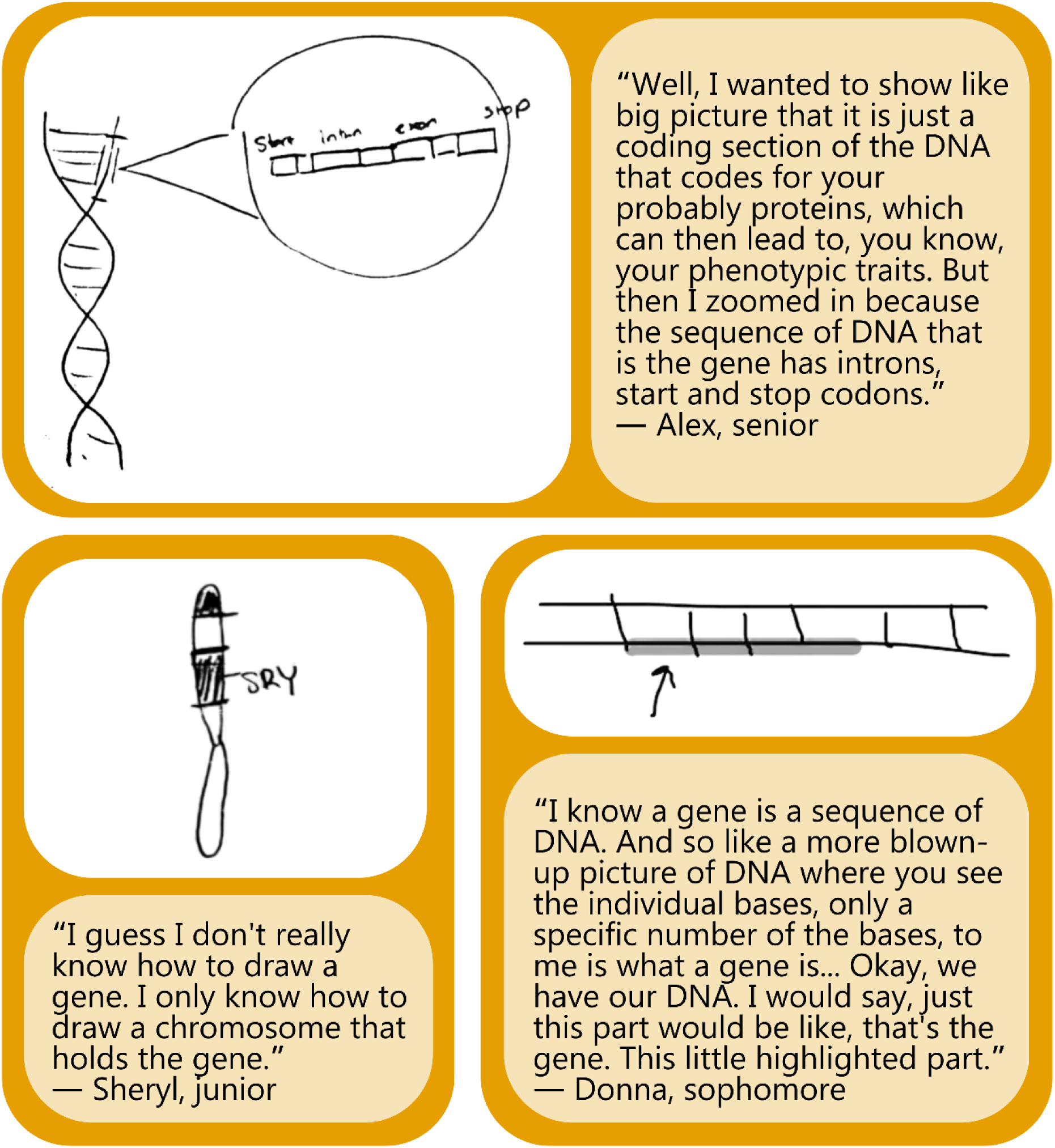
Example sketches that contain scale errors. Each sketch here shows an error in representing the scale of a gene. Alex and Donna show a gene as corresponding to only a few base pairs, while Sheryl shows a single gene taking up a large portion of a chromosome.

### What did errors in abstraction look like?

Many of the errors in abstraction occurred in the highly-abstract representations of chromosomes and genes. We have an extensive discussion of the common errors in drawing and interpreting highly-abstract X-shaped chromosomes and Punnett squares (which are included in the highly-abstract gene name location on the DNA Landscape) in our previous research (10), so we focus on abstraction errors in other locations on the DNA Landscape in Figure 4.

**Figure 4.**
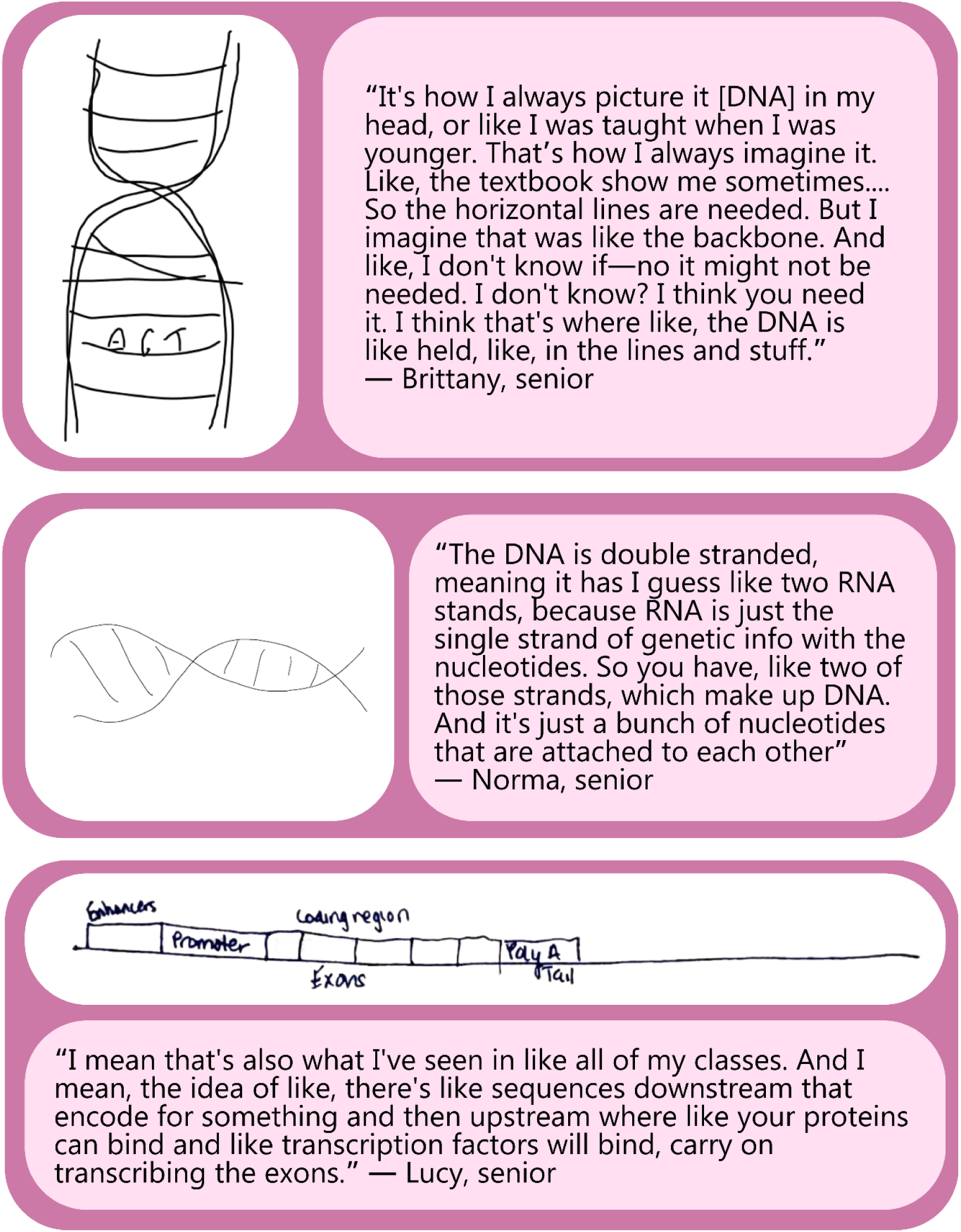
Example sketches that contain abstraction errors. The sketches made by Brittany and Norma indicate misunderstandings of the conventional meaning of the double helix shape commonly used to represent DNA. Lucy’s sketch indicates a misunderstanding of the conventions for representing genes with a box and line diagram.

We were surprised to see how many different ways students were misinterpreting the conventional double-helix shape of DNA, and we include two examples here. Brittany interpreted the phrase “double helix” to mean that there were two strands on each side of the structure. When completing her sketch, she used two pen strokes on each side of the twisted X-shape to emphasize the “double” nature of the double helix. Brittany’s drawing also indicated a misunderstanding of the phrase “backbone” that is conventionally used to refer to the alternating deoxyribose sugars and phosphate groups in a DNA strand. In her sketch, Brittany referred to the horizontal rung-shaped lines as the backbone of DNA and described this horizontal backbone as where DNA “is held.” She wrote out a series of three letters typically associated with nucleobases that were “held” upon the horizontal lines as if upon a shelf.

In comparison, Norma’s sketch of DNA is much more conventional in appearance. At first glance, the double-helix shape looks like what an expert might quickly draw to illustrate the double-helix shape of DNA. However, when Norma described her sketch, we can see that she has misunderstood the conventions — her double-stranded helix was actually representing “two RNA strands.” While Norma replicated a conventional drawing of DNA, she had an unconventional understanding of how experts typically use the double-helix shape to convey meaning about the two-stranded structure of DNA.

We also found several errors in the way students think about the meaning of conventional box-and-line diagrams of genes. Lucy’s gene here is entirely composed of boxes — the enhancer, promoter, exons, and poly A tail in the sketch are all represented with boxes. The line in Lucy’s box-and-line is only represented here after the end of the sequence of boxes has ended, which contrasts with the more conventional use of lines to illustrate untranscribed regions in genes. Lucy also mixes conventions of representing DNA and mRNA. Note how the “coding regions” of the gene consisting exclusively of exons, as if the gene had already undergone splicing. The gene in this sketch also resembles mRNA in its inclusion of a poly A tail. Despite multiple misuses of the conventions, Lucy insisted that this type of diagram was what she had seen in “all of [her] classes.”

### What does unconventional abstraction look like?

Student sketches were unconventional when the way that a student depicted a nucleotide, gene, or chromosome was not necessarily incorrect but did not match the standard uses of shapes, symbols, or sometimes colors, that experts typically rely on to convey meaning about molecular biology topics. In Figure 5, we see that students often use the correct vocabulary to describe molecular structures yet their sketches leave remaining questions about whether they truly understand the vocabulary they used. Consider the two chromatids in Sylvia’s sketch of a chromosome. Sylvia correctly states that chromosomes are made of two chromatids, yet she only colored in one of the chromatids in the pair. This color coding is unconventional because experts often use different colors to visually distinguish between non-homologous chromosomes in diagrams of mitosis or in karyotypes. Sister chromatids contain genetically identical information, so coloring only one of the chromatids conventionally signals a difference between the two. While Sylvia uses the correct words, we do not know if she robustly understands chromosome structure based on her drawing.

**Figure 5.**
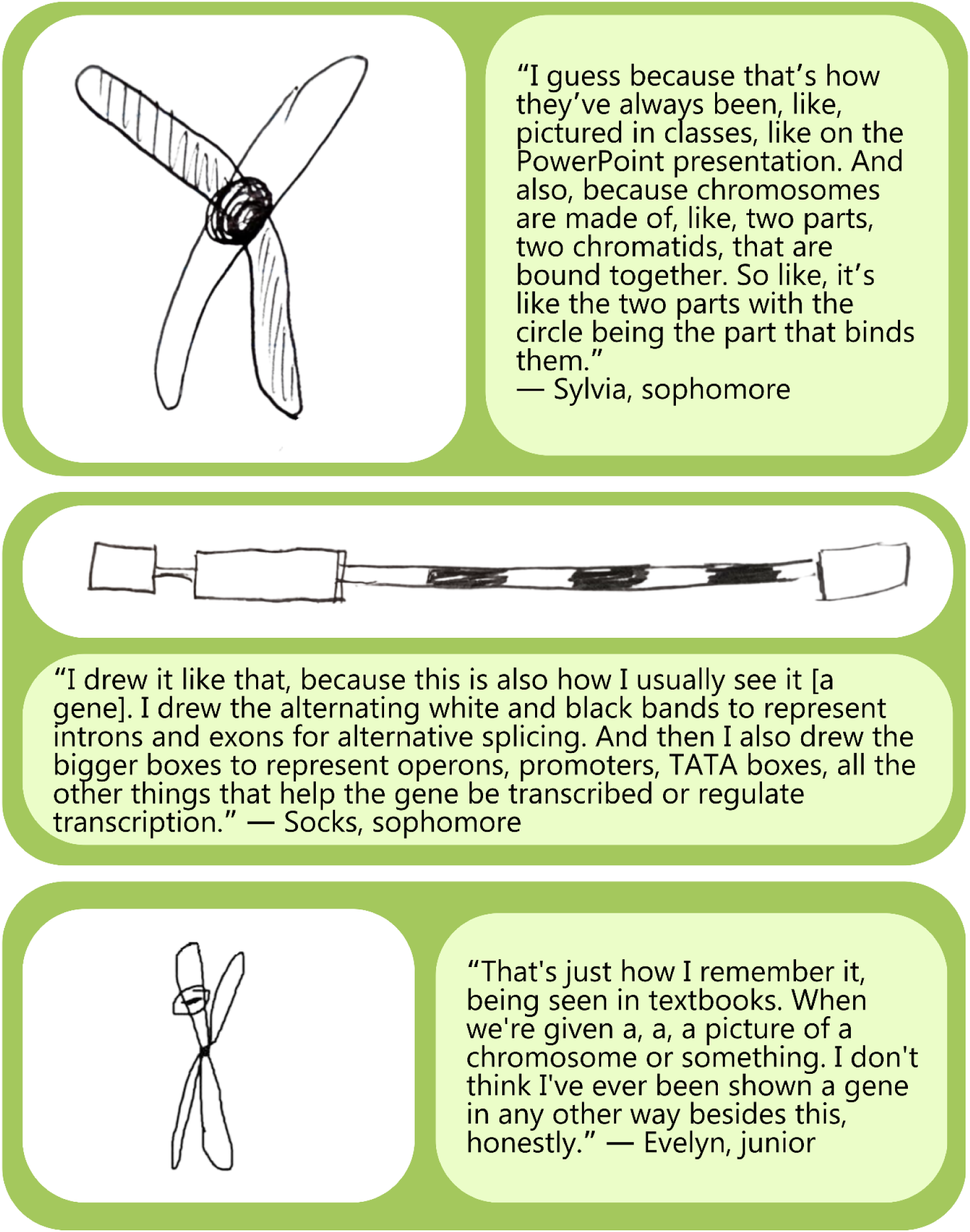
Example sketches that contain unconventional abstraction. Sylvia’s sketch uses unconventional color coding in a representation of a replicated chromosome. Socks unconventionally ascribes meaning and colors to the conventional shapes of a box and line diagram of a gene. Evelyn’s sketch is unconventional in that a gene is only indicated on one chromatid in a representation of a replicated chromosome.

We see a similar situation where correct words are paired with unclear symbolism in Socks’ box-and-line sketch of a gene. Here, the boxes are not conventionally representing exons, but instead are supposed to be operons, promoters, and TATA boxes. This sketch also uses the narrow line to represent both introns and exons, distinguishing the two by unconventional color coding. While someone with more expertise might use similar words and similar shapes to describe a gene, the labels and color coding that Socks uses to explain the sketch leave questions about the extent to which they understand gene structure.

In a third case, Evelyn represents a gene as a narrow band on a chromosome, which could be a fairly conventional representation of genes. However, the chromosome is replicated, and the gene is only indicated on one of two arms. We do not know from this sketch or from her description whether Evelyn understands that the same gene would also be present in the same location on other sister chromatid.

### What does it look like when sketches contain both errors in scale and abstraction?

While only a small portion of sketches contained both errors in scale and abstraction (n = 14), the types of errors we observed in these sketches indicate deep misunderstandings of foundational structures in molecular biology. These sketches incorrectly mixed and matched the conventions across locations on the DNA Landscape. We describe three such instances of students misusing conventions for scale and abstraction in Figure 6.

**Figure 6.**
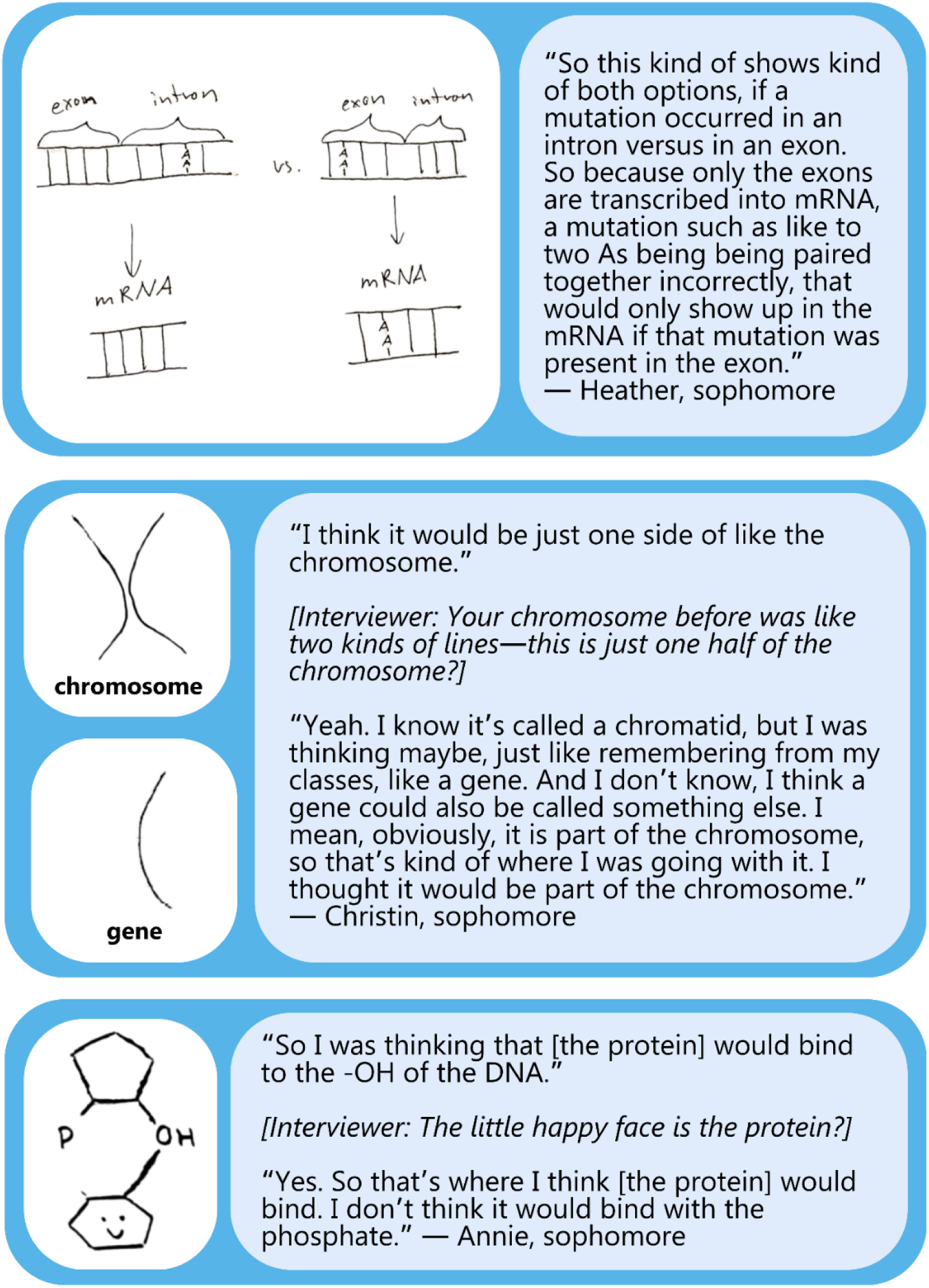
Example sketches that contain both errors in scale and abstraction. Heather’s sketch misrepresents the scale of a gene and misuses the conventions of the ladder shape to represent mRNA. When compared to her original sketch of a chromosome, Christin’s sketch of a gene indicates a misunderstanding of the size and structure of a gene. Text labels were added to Christin’s sketches for clarity. Annie’s sketch indicates a misunderstanding of the size and structure of both nucleotides and proteins.

Similar to Donna (Figure 3), Heather misused the conventions of the nucleotide ladder to represent exons and introns as structures consisting of approximately 3 nucleotide base pairs. Heather drew this sketch as a representation of the true-false statement in GenBio-MAPS: “Only mutations that occur in the exons can have an impact on the cell,” and chose to represent the adenine-adenine pairing as a mutation that might occur. Heather correctly transcribed DNA into mRNA, but continued to use the conventions of the ladder to represent mRNA as a double-stranded structure.

Christin’s sketch of a gene indicated a misunderstanding of both the conventions of abstract X-shaped chromosomes as well as the scale of a gene. In the first part of the interview, Christin originally drew a chromosome as two side-by-side lines. When we asked Christin to draw a gene later in the interview, she drew a gene using the same shape, size, and length as just one of the lines in the original chromosome sketch. She verbally indicated that her sketch showed that a single gene was the entire length of a chromosome. While both genes and chromosomes can be represented with lines, Christin misunderstood that the similarly-shaped lines representing genes and chromosomes conventionally indicate vastly different scales of genetic information.

Annie was among many of the participants who substantially misrepresented the scale of nucleotides and proteins. Here, Annie represents a protein as a smiling hexagon that is binding to the –OH group of a single DNA nucleotide. In addition to representing an entire protein at the same scale as a single nucleotide, Annie makes an interesting choice to use a hexagon to represent the protein structure. While proteins can be abstractly represented using a variety of shapes, ranging from circles to amorphous blobs, experts rarely use hexagons to represent proteins because hexagons are conventionally used to represent sugars.

### What are students doing to accurately convey scale?

We were encouraged to see that some students were cognizant of the limitations of accurately conveying scale in a quick sketch. Several students acknowledged how they would like to improve their sketch to be more realistic. For example, Piper drew a short sequence of letters to represent a gene and used ellipsis and the label “# of bps [base pairs] long” to signal that genes are much longer than what is feasible to represent in a sketch (Figure 7). While students like Piper demonstrated their understanding of scale by pointing out the shortfalls in their sketches, this was not common across all students. Unless the student verbally or visually indicated a limitation of their sketch, we assumed that students’ sketches were representative of their mental models of scale.

**Figure 7.**
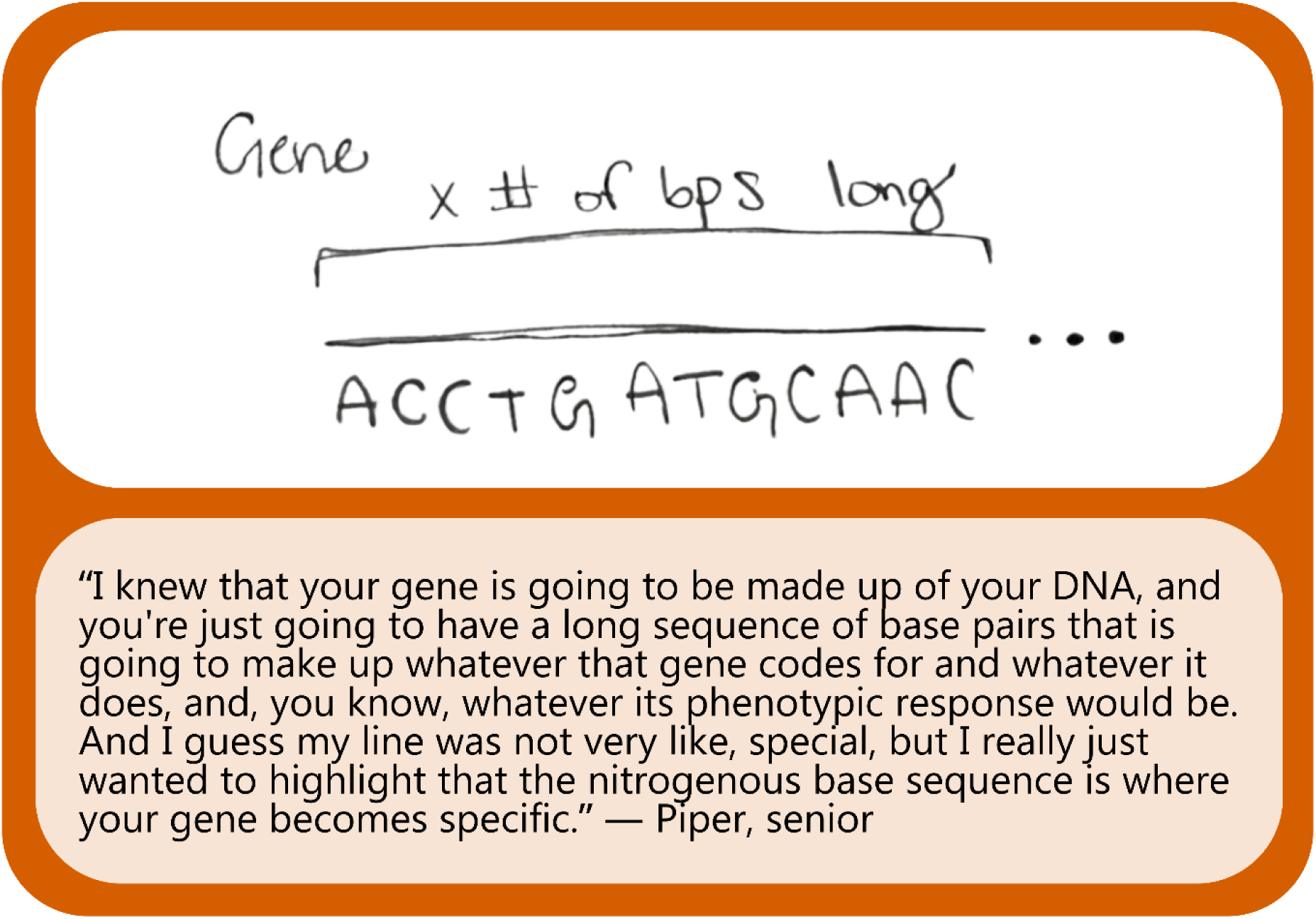
Example sketch that is annotated in a way that reflects an understanding of scale. Piper annotated the short nucleotide sequence with an ellipsis and the label “X # of [base pairs] long,” which indicates she understood that genes are longer sequences than what is practical to draw in a quick sketch.

## Discussion

We analyzed student sketches of nucleotides, genes, and chromosomes and found that most students misused or misunderstood the visual conventions for scale and abstraction that experts typically use to represent these molecular structures. Some of the most recognizable symbols in biology — the double helix and the X-shaped chromosome — were among the most commonly misunderstood. Understanding scale and abstraction in molecular structures is foundational for visual literacy (6, 13), and our findings suggest that undergraduate students may need additional instructional supports to develop fluent visual literacy in molecular biology.

### Supporting student understanding of conventions for scale

Scale is so important to science that it is included as a *crosscutting concept* in a widely-used framework for science education (20), yet scale is rarely emphasized or assessed in undergraduate biology courses (21). To help better incorporate scale into biology teaching and learning, we recommend using appropriately scaled physical models of molecular structures to provide a foundation for student thinking (22). When using physical models in a classroom may be limited by monetary or class size constraints, making analogies to familiar objects can also be a fruitful way to build connections in student understanding of molecular scale (23). For example, when discussing the size of a gene or the size of a chromosome, instructors can make a comparison to a skein of yarn. Instructors can challenge students to determine how many inches a gene would be if nucleotides are scaled to the size of a single strand of yarn, and to consider the relationship between the wrapped nature of a skein and the condensed nature of a chromosome. Relating nucleotides, genes, and chromosomes to a physical object like yarn may help students visualize that a gene can be tens of thousands of nucleotides, and might help navigate students away from misunderstandings like a single gene comprises a substantial portion (or the entire length) of a chromosome. After using yarn as an example, we suggest instructors ask students to translate that analogy back to the typical conventions for representing scale in molecular biology. Instructors can show various examples of sample sketches in which genes are represented at conventional and unconventional sizes and ask students to discuss in groups and explain which representation is most accurately portrays the scale of a gene.

Instructors may also consider asking their class to evaluate each other’s drawings of scale. In this study, we asked students to “zoom in” and “zoom out” of molecular structures, and these prompts generated some of the most interesting misrepresentations of scale in student sketches. Instructors can provide students with drawing materials in class, ask students to “zoom in” and “zoom out” from nucleotides, genes, and/or chromosomes, and then ask students to evaluate each other’s sketches. In such an approach, we suggest instructors end the activity with their own expert analysis of student sketches to positively reinforce which conventions were used correctly and steer students away from incorrect representations of scale.

### Supporting student understanding of conventions in abstract representations of DNA

Biologists have multiple ways of visually representing the same molecular structures, but these multiple representations can pose challenges for students. While experts can fluently translate between representations of nucleotides as letters, as ladders, and as chemical structures because their experience using such symbols has conferred “representational competence” (24), students may need additional guidance to correctly choose which representation is most appropriate to communicate a certain point. To help students develop this representational competence, instructors can be explicit about how biologists use abstraction to focus viewers on the most salient features of the structure. Instructors may ask students to consider why a biologist might use letters rather than chemical structures when the most important information to convey to another biologist is the sequence of nucleobases. To help students understand why there are multiple levels of abstraction for representing the same molecular structures, we recommend using active-learning opportunities in class where students discuss in small groups why and when a biologist might use certain representations to communicate. Instructors may also consider incorporating “figure analysis” activities in which students work in groups to examine and discuss the meaning and the limitations of the abstract symbolism in textbook figures (25).

In addition to discussing polished textbook figures, instructors may find it a useful and insightful learning task to have students evaluate unconventional representations of DNA. Instructors may draw their own intentionally unconventional sketches, find examples of unconventional representations published online in image banks such as Adobe Stock, or create unconventional representations using artificial intelligence. Asking students to consider the strengths and limitations of a representation and how the representation conventionally or unconventionally portrays molecular structures may help students understand what features in abstractions enable effective communication between biologists.

### Using the DNA Landscape to support student understanding of scale and abstraction

Our findings suggest that most students may need support to understand and use visual conventions for representing DNA. We recommend instructors use the DNA Landscape (12) as an instructional tool to help scaffold their teaching about visual representations of DNA. When designing a lesson, a unit, or a course, instructors should consider how they are incorporating representations of DNA from locations on the DNA Landscape. Instructors can explicitly compare and contrast representations within the same column, highlighting how similar meaning about a structure is conveyed at different levels of abstraction. We also recommend that instructors consult the DNA Landscape when they are using representations that span across columns, as spanning columns within a representation can muddle scalar relationships. For example, we saw that many students were representing entire genes (from the middle column of the DNA Landscape) with letters (from the left column of the DNA Landscape), and they typically represented entire genes with fewer than 10 letters. Mixing and matching across columns can create confusion about scale. If using letters to represent a gene, instructors may consider following Piper’s example in Figure 7 and explicitly writing out an estimate for the number of letters that might compose the entire gene.

If instructors emphasize visual literacy skills in their teaching, they should make sure that these priorities are also reflected in their assessments. Despite the frequency with which visual models are used in teaching, we previously found that visual models are largely absent from undergraduate biology exams (26). If instructors are teaching students using representations from across the DNA Landscape, we encourage instructors to also assess students on their understanding of representations from across the DNA Landscape. In this present study, we found that drawings were an effective way to quickly assess student understanding of molecular structures, but we were only able to glean many of these understandings by comparing both sketches and verbal descriptions (such as in Norma’s double-stranded helix representing two RNA strands in Figure 4). We recommend that instructors consult the Three-Dimensional Learning Assessment Protocol (27) for criteria that may guide the creation of assessment questions that can engage students in productively reasoning about visual models.

### Limitations

We necessarily focused our analysis on student errors and misunderstandings to sharpen our focus on the areas of undergraduate biology instruction that may need additional scaffolding and support. This decision was not meant to undervalue students’ productive thinking about molecular biology.

Our sample consisted of students from two institutions, and such, the types of errors in scale and abstraction that we observed may not be representative of student thinking in other populations.

## Conclusion

Biologists rely on conventional visual models to communicate about molecular structures, yet the conventions in such models are often misunderstood by undergraduate biology students. Even though the shapes and symbols to represent nucleotides, genes, and chromosomes are ubiquitous in undergraduate biology courses, students may not be interpreting the meaning of the shapes and symbols in the same ways as experts. By asking students to draw molecular structures, we saw that many students had sketchy understandings of foundational biology concepts. As visual literacy is central for communicating about molecular biology, we recommend instructors closely examine the visual representations they use in their courses and consider the ways in which they can reinforce student understanding of the conventions for representing scale and abstraction.

## Acknowledgements

We thank the students who participated in this research. We thank Kerstyn Gay for creating the illustrations in the DNA Landscape in Figure 1. Funding provided by NSF DGE 2222337. Any opinions, findings, and conclusions or recommendations expressed in this material are those of the author(s) and do not necessarily reflect the views of the National Science Foundation.

